# A novel lineage of *Chilomastix mesnili* in wild Moroccan Barbary macaques (*Macaca sylvanus*)

**DOI:** 10.64898/2026.10.27.762143

**Authors:** Allegra V. Brady, Michael Jarman, Mohamed Arbaoui, Liz Campbell, Sidi I. Cherkaoui, Majda El-Bouhamdi, Matthew Gardiner, Sophie Hughes, Joe Linscott, Nicola F. Koyama, Mohamed Naimi, Mirko Pegoraro, Peter Shum, Patrick J. Tkaczynski, Gareth D. Weedall

## Abstract

*Chilomastix* spp. are flagellated protists found in the intestines of a broad range of hosts. They are generally non-pathogenic and have been little studied, especially using molecular techniques, and significant taxonomic uncertainty regarding the host specificity of lineages remains. As part of a study of intestinal microbial communities of wild Barbary macaques (*Macaca sylvanus*) from Morocco, we mined 50 faecal metagenomic sequence libraries and found 11 rRNA sequences matching *Chilomastix*. Compared to published 18S rRNA sequences, the Barbary macaque *Chilomastix* sequences formed a well-supported clade distinct from human-derived *C. mesnili* ST1 from Indonesia. The macaque *Chilomastix* sequences cluster with a single pig-derived isolate and two sequences from Japanese macaques (*Macaca fuscata*). This indicates that *M. sylvanus* harbours a lineage of *Chilomastix* distinct from the common human strain of *C. mesnili* and geographically widespread. We were also able to reconstruct the complete rDNA operon from this lineage.

## MAIN TEXT

Protozoa exhibit a broad spectrum of host specificity, ranging from specialists adapted to a single host species to generalists capable of infecting many species. Species identification based primarily on morphological characteristics can lead to misclassification due to the presence of cryptic species, phenotypic plasticity, and convergent evolution, whereas genetically distinct organisms exhibit similar structural features (**Ko *et al*., 2013**). Determining host specificity in intestinal protozoa is complicated due to the limitations of morphology-based identification (**Chabé *et al*., 2017**). Molecular techniques have shown that host origin may reflect underlying genetic divergence, suggesting that morphologically identical isolates from different hosts could represent distinct, host-associated species (**Jiang *et al*., 2025**).

*Chilomastix* spp. are flagellated protists found in the intestines of both vertebrates and invertebrates and also free-living (**Cevasco, 2022; Jiang *et al*., 2025**). *Chilomastix mesnili*, the best characterized species in the genus, is most frequently detected in humans (**Suzuki et *al*., 2023; Jiang *et al*., 2025**). It is generally regarded as a harmless commensal, adapted to low-oxygen environments within the large intestine (**Kulda *et al*., 2017**), though it has been reported as an opportunistic pathogen in immunocompromised individuals (**Morimoto *et al*., 1996; Kaniyarakkal *et al*., 2016**). *C. mesnili* has a simple life cycle, without intermediate hosts or vectors: motile trophozoites live in the host gut, and resistant cysts passed in faeces are transmitted to new hosts via ingestion, excystation taking place in the intestinal lumen of the new host (**Leiva, 1921**). Due to its well characterised morphology, life cycle, and host association, *C. mesnili* is commonly used as a reference species for comparison with *Chilomastix* isolates identified in other vertebrate hosts (**Suzuki et al., 2023**). More than 30 species have been described in the genus, based on morphology and host species, but not on genetic information (**Kulda *et al*., 2017**). The few available studies of genetic diversity in the genus suggest a very high level of genetic divergence among putative species (**Jiang *et al*., 2025**).

The largest study of molecular diversity of *Chilomastix* to date was by Jiang *et al*. (**2025**). In this work, humans and other domestic and peridomestic animals (ducks, chickens, water buffalo and rats) were sampled from a single location (Wainyapu village, Sumba Island, Indonesia). A high level of genetic diversity was reported among *Chilomastix* partial 18S rRNA gene sequences, which grouped into lineages likely representing host-associated species. Rat- and chicken-derived lineages were highly divergent from pig- and human-derived lineages (which likely represent *C. mesnili*). They found a rat-derived lineage (‘*Chilomastix bettencourti*-like’) and a chicken-derived lineage (‘*Chilomastix gallinarium*-like’), both highly divergent from *C. mesnili*, of which they found two distinct lineages (Sub-Type 1, ST1, and ST2). ST2 was further subdivided into ST2-1 (predominantly human hosts) and ST2-2 (predominantly non-human hosts). ST1, conversely, was found mostly in human hosts (aside from a single pig host).

As part of a larger study of intestinal microbial communities of wild Barbary macaques (*Macaca sylvanus*) in Morocco, stool samples of 50 individuals were used for shotgun metagenomic sequencing. DNA isolation was automated on a KingFisher Apex instrument using the MagMAX Microbiome Ultra Nucleic Acid Isolation Kit. Library preparation and sequencing were performed on an Illumina NovaSeq 6000 at the Centre for Genomics Research, University of Liverpool. We identified the presence of *Chilomastix* in this molecular sequence data and studied its relationship to published genetic data for the genus. Many populations of Barbary macaques, particularly those in Gibraltar and high-traffic areas of North Africa, live near human settlements (**Kwiatt, 2017**) and/or come into close contact with humans at ecotourism sites (**Waterman *et al*., 2000**). This anthropogenic overlap creates a potential pathway for cross-species transmission of generalist parasites (**Borremans *et al*., 2019**), requiring a deeper understanding of the parasites they harbour.

From 50 Barbary macaque (*Macaca sylvanus*) faecal samples screened, eleven putative *Chilomastix* sequences were identified (from eleven different macaques), based on full or partial match to an 18S SSU rRNA query sequence. Of these 11 sequences (submitted to GenBank: accessions QB174119-QB174129), five sequences fully spanning the central part of the 18S SSU rRNA gene were aligned to an additional 24 *Chilomastix mesnili* 18S SSU rRNA gene sequences from the GenBank database (**Table 1**), using MUSCLE (**Edgar, 2004**) with default settings, implemented in SeaView v5 (**Gouy *et al*., 2021**). The alignment was used to construct a neighbour-joining phylogeny with 1000 bootstrap replicates using BioNJ (**Gascuel, 1997**) with the Kimura 2-parameter distance correction model, implemented in SeaView v5 (**Gouy *et al*., 2021**). A maximum likelihood tree for the same alignment produced the same tree topology (data not shown). In the phylogeny (**Figure 1**) all sequences derived from Barbary and Japanese macaques (*Macaca fuscata*) (**Suzuki *et al*., 2023**) clustered together with a single pig-derived sequence (**Jiang *et al*., 2025**). Jiang *et al*. (**2025**) included this pig-derived sequence among ST1 as it was a single isolate, despite a relatively long branch separating it from the other ST1 sequences. Our data support including it alongside the macaque-derived sequences to form a separate, non-human-host-specific clade (which we call ST1-3). We also suggest, based on branch lengths, that the human-derived ST1 sequences could be divided into sub-groups ST1-1 and ST1-2 (**Figure 1**). ST1-1 contains human-derived *Chilomastix mesnili* all from one site in Indonesia (**Jiang *et al*., 2025**), while ST1-2 were collected from a person returning from South America to the Czech Republic (**Cepicka *et al*., 2008**). Therefore, it is not possible with this data to disentangle the effect of geographical population structure from species divergence. Greater sampling from more geographical locations will help to disentangle these effects. By contrast, our proposed ST1-3 contains sequences from three widely geographically separated locations and three host species. Despite this, the sequences are all closely related to one another, suggesting a single clade that is widely distributed and infects more than one mammalian host.

**Table 1.** *Chilomastix mesnili* 18S SSU rRNA gene sequences used in this study.

| Accession | Taxon | Host | Length (bp) | Reference |
| --- | --- | --- | --- | --- |
| LC789661 | *C. mesnili* ST1-1 | human | 1389 | Jiang *et al.* (2025) |
| LC789662 | *C. mesnili* ST1-1 | human | 1387 | Jiang *et al.* (2025) |
| LC789717 | *C. mesnili* ST1-1 | human | 1389 | Jiang *et al.* (2025) |
| LC789684 | *C. mesnili* ST1-1 | human | 1386 | Jiang *et al.* (2025) |
| LC789668 | *C. mesnili* ST1-1 | human | 1389 | Jiang *et al.* (2025) |
| EU009464 | *C. mesnili* ST1-2 | human <sup>1</sup> | 2453 | Cepicka *et al.* (2008) |
| EU009466 | *C. mesnili* ST1-2 | human <sup>1</sup> | 2344 | Cepicka *et al.* (2008) |
| LC789686 | *C. mesnili* ST1-3 | pig | 1406 | Jiang *et al.* (2025) |
| LC752183 | *C. mesnili* ST1-3 | Japanese macaque | 878 | Suzuki *et al.* (2023) |
| LC771049 | *C. mesnili* ST1-3 | Japanese macaque | 878 | Suzuki *et al.* (2023) |
| BM_1160529_L13566_rc | *C. mesnili* ST1-3 | Barbary macaque | 13566 | This study |
| BM_27067_L13660_rc | *C. mesnili* ST1-3 | Barbary macaque | 13660 | This study |
| BM_10954_L13657 | *C. mesnili* ST1-3 | Barbary macaque | 13657 | This study |
| BM_3933_L4018_rc | *C. mesnili* ST1-3 | Barbary macaque | 4018 | This study |
| BM_385429_L13639_rc | *C. mesnili* ST1-3 | Barbary macaque | 13639 | This study |
| KC960587 | *C. mesnili* ST2-1 | human | 2447 | Cepika (Unpublished) |
| KC960588 | *C. mesnili* ST2-1 | human | 2445 | Cepika (Unpublished) |
| LC789665 | *C. mesnili* ST2-1 | human | 1435 | Jiang *et al.* (2025) |
| LC789666 | *C. mesnili* ST2-1 | human | 1441 | Jiang *et al.* (2025) |
| LC789673 | *C. mesnili* ST2-2 | pig | 1446 | Jiang *et al.* (2025) |
| LC789657 | *C. mesnili* ST2-2 | water buffalo | 1461 | Jiang *et al.* (2025) |
| LC789672 | *C. mesnili* ST2-2 | pig | 1461 | Jiang *et al.* (2025) |
| LC789720 | *C. mesnili* ST2-2 | human | 1459 | Jiang *et al.* (2025) |
| LC789676 | *C. mesnili* ST2-2 | pig | 1460 | Jiang *et al.* (2025) |
| LC789678 | *C. mesnili* ST2-2 | pig | 1460 | Jiang *et al.* (2025) |
| LC789687 | *C. mesnili* ST2-2 | pig | 1455 | Jiang *et al.* (2025) |
| LC789729 | *C. mesnili* ST2-2 | pig | 1459 | Jiang *et al.* (2025) |
| LC789675 | *C. mesnili* ST2-2 | pig | 1454 | Jiang *et al.* (2025) |
| LC789688 | *C. mesnili* ST2-2 | pig, chicken, dog | 1453 | Jiang *et al.* (2025) |

<sup>1</sup> Both sequences were derived from a single human, a traveller returning to the Czech Republic from South America.

**Figure 1.**
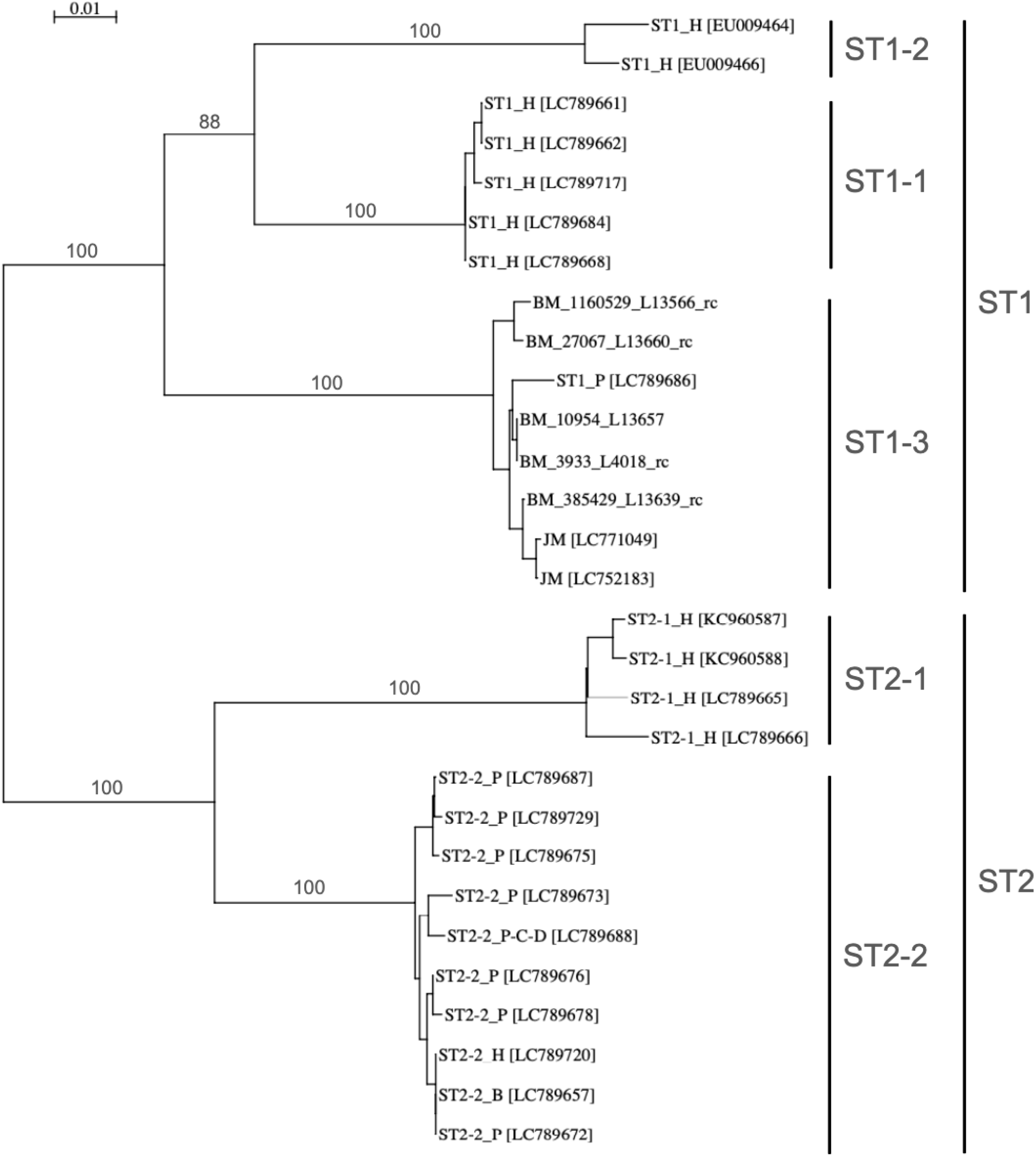
Phylogeny of an approximately 800 bp region of the 18S SSU rRNA gene of *Chilomastix mesnili*. Sequence subtypes (ST) designated by Jiang *et al*. (2025) are shown in the taxon names (ST1, ST2-1, ST2-2). These and new proposed sequence types (ST1-1, ST1-2, ST1-3) are also indicated by vertical bars on the right. In taxon names, host species are indicated by capital letters (H=human, P=pig, B=buffalo, C=chicken, D=dog, JM=Japanese macaque, BM=Barbary macaque). Percentages of 1000 bootstrap replicates supporting major branches are shown (they are not shown for the shorter branches for clarity). The phylogeny was constructed using the BioNJ method with 1000 bootstrap replicates and using Kimura’s 2-parameter distance correction model.

To date, the longest DNA sequence determined from any *Chilomastix* species is an approximately 2.5 kb region of the 18S SSU rRNA gene. Here, we used our *de novo* assembled data to reconstruct the complete rDNA amplicon of *C. mesnili* ST1-3. Four of our assembled contigs were similar in length: 13,566-13,660 bp. We annotated 18S, 5.8S and 28S rRNA genes in these contigs using Infernal cmscan (INFERence of RNA ALignments Covariance Model scan; **Nawrocki and Eddy, 2013**) via the EMBL-EBI Job Dispatcher (**Madeira *et al*., 2004**). We manually edited the sequences to harmonise gene order and aligned them.

Consistency in sequence similarity among independently assembled sequences from separate macaques indicates that they are not mis-assemblies. The results show that the rDNA operon consists of 18S (2494-5 bp), ITS1 (400 bp), 5.8S (140 bp), ITS2 (564-568 bp), 28S (5696-5700-5704 bp) and an intergenic spacer (IGS; 4209-4222-4224 bp) between tandemly repeating units. The 28S rRNA gene appears to contain a 788 bp group I intron (**Figure 2**). While *C. mesnili* 18S SSU rRNA gene size is larger than other species (mouse ∼1.9 kb, *Drosophila* and yeast ∼2.0 kb, *Xenopus* ∼1.8 kb, *Giardia* ∼1.4 kb), the 28S LSU rRNA gene size (when the 788 bp intron is removed) is of a similar length to the human (∼5 kb) and mouse (∼4.7 kb) orthologues, larger than *Drosophila* and yeast (∼4.1 and ∼3.8 kb, respectively) and much larger than *Giardia* (∼2.7 kb).

**Figure 2.**
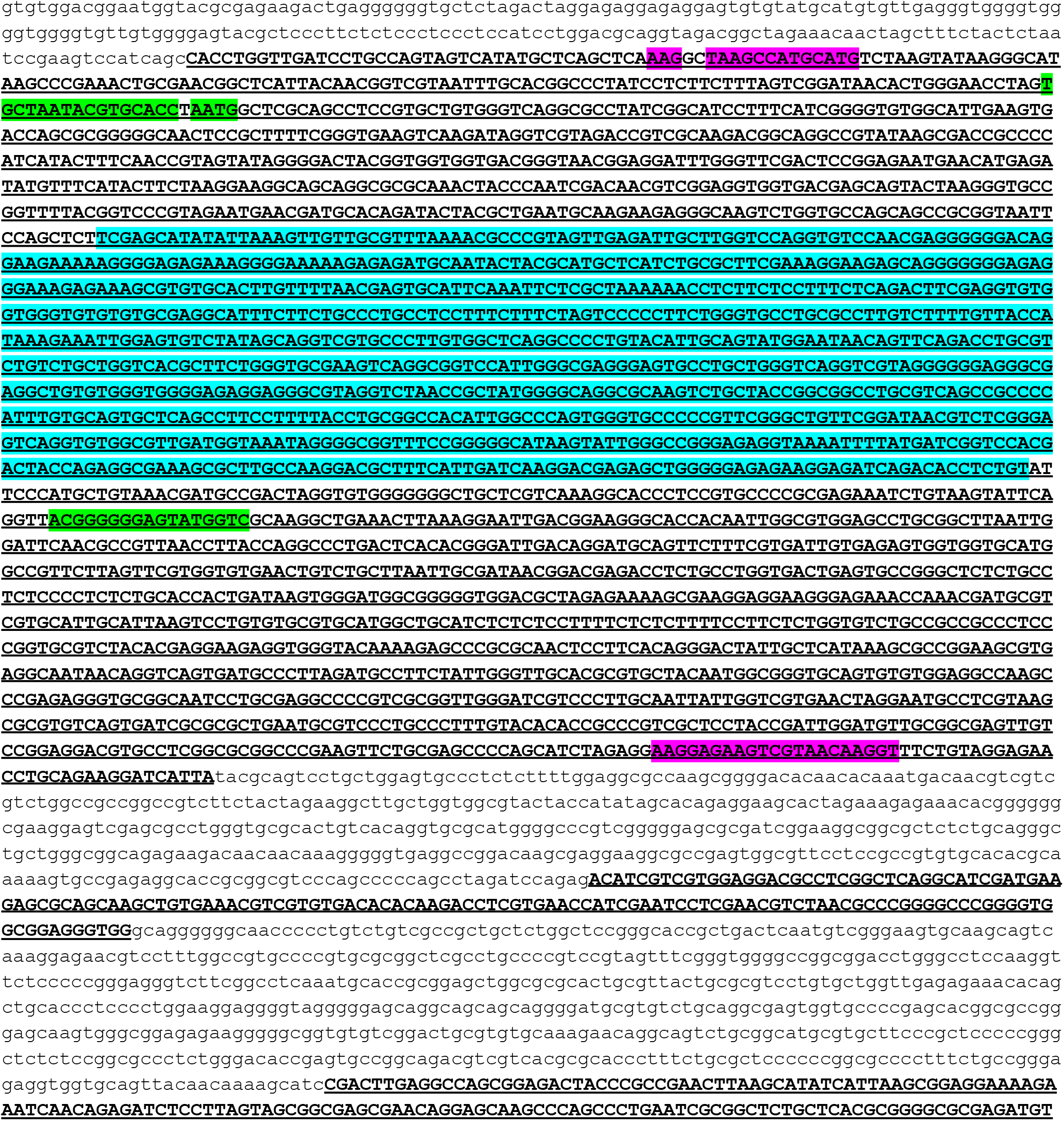
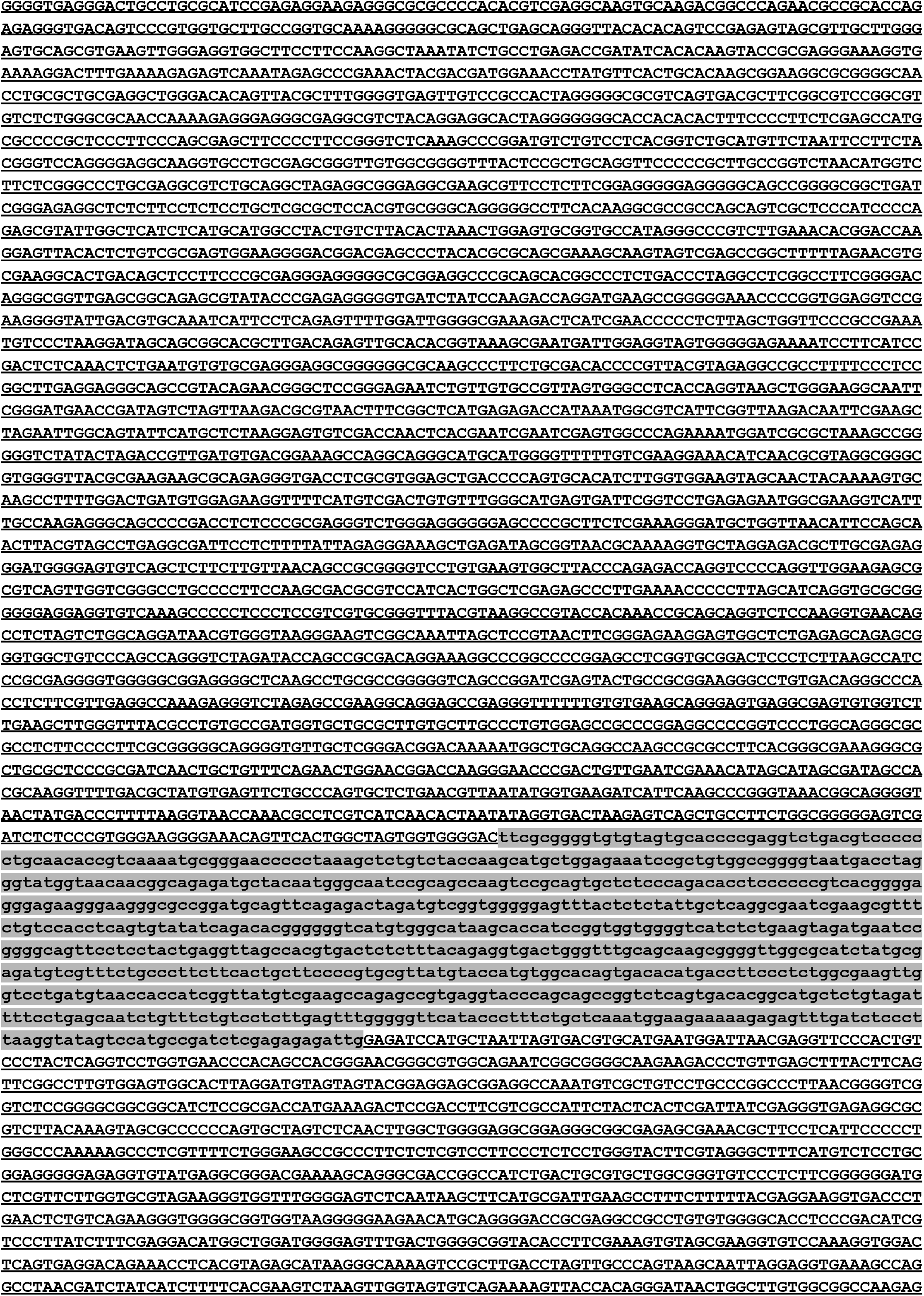
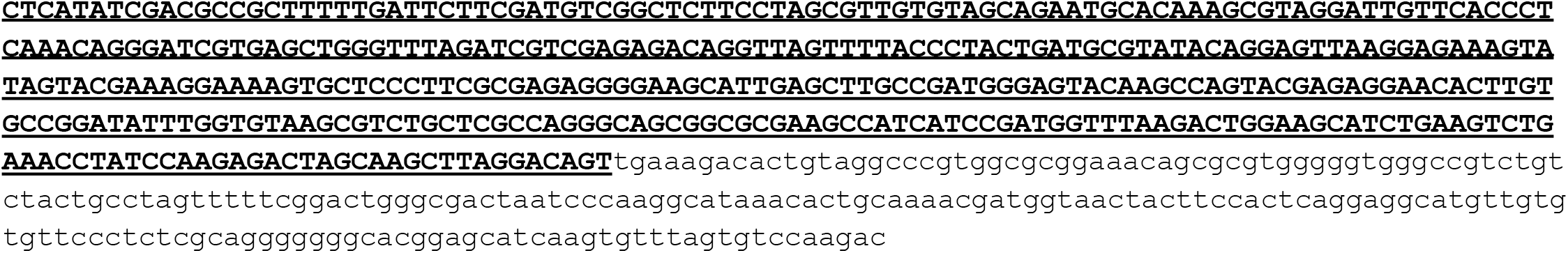
Representative sequence of the ribosomal DNA operon, not including the full intergenic spacer (IGS) of approximately 4 kb. The sequence shown is taken from the sequence BM_10954_L13657 generated in this study. 200 bp upstream of the start of the 18S SSU rRNA gene and 200 bp downstream of the end of the 28S LSU rRNA gene are shown in lower case, as are the 400 bp internal transcribed spacer ITS1, the 568 bp ITS2 and a 788 bp putative intron in the 28S LSU rRNA gene. 18S (2495 bp), 5.8S (140 bp) and 28S (5704 bp) rRNA genes are in bold and underlined. A predicted group I catalytic intron (788 bp) in the 28S LSU rRNA gene is highlighted in grey and shown in lower case. Within the 18S SSU rRNA gene the positions of the outer (pink) and inner (green) nested primer pairs used by Jiang *et al*. (**2025**) are indicated (mismatched nucleotides are un-highlighted). The region we used for the multiple sequence alignment and phylogenetic reconstruction, determined by the length of two *Chilomastix* sequences from Japanese macaques (**Suzuki *et al*., 2023**) is shaded in cyan.

To our knowledge, *Giardia* spp. are the closest relatives to *Chilomastix* spp. for which complete sequences of the rDNA operon have been determined. In this genus the operon is short compared to other eukaryotes, with both the 18S SSU and 28S LSU genes being considerably shorter (**Edlind and Chakraborty, 1987**). By contrast, *Chilomastix* is more typical of other eukaryotes. The intergenic spacer (IGS) between the end of the 28S LSU rRNA gene and the beginning of the 18S rRNA SSU rRNA gene of the next rDNA repeat was approximately 4.2 kb in these putative *Chilomastix mesnili* sequences. This is much shorter than those measured in a small number of vertebrates, in which the IGS ranged from 8-37 kb (**Macdonald *et al*., 2024**).

Among the sequences not used in the alignment and phylogeny due to incomplete coverage of the central part of the 18S SSU rRNA gene, one sequence, BM_969175_L8964, differed substantially from the others, presenting shorter ITS1 (342 bp), ITS2 (473 bp) and 28S LSU rRNA gene (4764 bp) and aligning poorly across the IGS region. Although the BLAST of this sequence against GenBank data returned *C. mesnili* as the highest scoring match, it is possible that it represents another retortamonad species with no representative rDNA data in GenBank.

## ACKNOWLEDGEMENTS

We thank Morocco’s L’Agence Nationale des Eaux et Forêts for permission to collect the samples and data. Work was funded by a NERC Urgency grant (NE/Y00647X/1) and NEOF support (project # NEOF1726).

